# Gastruloids generated without exogenous Wnt activation develop anterior neural tissues

**DOI:** 10.1101/2020.10.10.334326

**Authors:** Mehmet U. Girgin, Matthias P. Lutolf

## Abstract

When stimulated with a pulse from an exogenous WNT pathway activator, small aggregates of mouse embryonic stem cells (ESCs) can undergo embryo-like axial morphogenesis and patterning along the three major body axes. However, these structures, called gastruloids, currently lack the anterior embryonic regions, such as those belonging to the brain. Here we describe an approach to generate gastruloids that have a more complete antero-posterior development. We used bioengineered hydrogel microwell arrays to promote the robust derivation of mouse ESCs into post-implantation epiblast-like (EPI) aggregates in a reproducible and scalable manner. These EPI aggregates break symmetry and axially elongate without external chemical stimulation. Inhibition of WNT signaling in early stages of development leads to the formation of gastruloids with anterior neural tissues. Thus, we provide a new tool to study the development of the mouse after implantation *in vitro*, especially the formation of anterior neural regions.

## Introduction

The development of the mouse embryo is an incredibly complicated and fascinating process that leads to a complete body plan with perfectly organized organs and tissues within just a few weeks. Numerous *in vivo* studies have been made to understand how this process takes place. Compared to other model organisms, however, the performance of genetic experiments on mice is often time-consuming and is complicated by the limited accessibility of the embryos after implantation. This has motivated the establishment of *in vitro* models to study early embryonic development.

The first attempts to model the embryonic development of the mouse *in vitro* began with an incidental observation. The analysis of teratocarcinomas from neonatal mouse testes showed a striking structural similarity to the developing mouse embryo (Stevens 1959). Embryonic carcinoma cells (EC) isolated from these tumors (Kleinsmith & Pierce 1964) were aggregated *in vitro* to form so-called “embryoid bodies” (EBs) which recapitulated aspects of the spatial organization of the endodermal and ectodermal layers of mouse embryos on day 5 of development (Martin & Evans 1975a; Martin & Evans 1975b). EC cells in EBs were then rapidly replaced by embryonic stem cells (ESCs) to generate cell types derived from all three germ layers (Bain et al. 1995; Kubo et al. 2004; Boheler et al. 2002). However, although countless important studies with embryoid bodies as model system have been performed, revealing the remarkable self-organizing potential of ESCs, including antero-posterior axis determination (Berge et al. 2008) and gastrulation-like events (Marikawa et al. 2009; Li et al. 2015; Samal et al. 2020), the lack of robustness in development and morphogenesis of EBs has made it difficult to study more complex aspects of embryonic development. An improved version of EBs, termed gastruloids, can be generated from smaller ESC aggregates, well-defined media and timed chemical stimulation, resulting in a stereotypical *in vivo*-like axial elongation and patterning (van den Brink et al. 2014; Turner et al. 2017; Beccari et al. 2018). These multicellular aggregates were shown to recapitulate spatio-temporal activation of *Hox* genes, a phenomenon that is evolutionarily conserved among vertebrates (Mallo et. al. 2013). Compared to the embryo, however, the development of the gastruloid is limited to the post-occipital region (Beccari et al. 2018), with anterior nervous tissues, which correspond to the forebrain, midbrain and hindbrain, largely absent.

The derivation of mouse gastruloids is based on the treatment of mouse ESC aggregates with the WNT agonist CHIR99021, which induces symmetry breaking and axial elongation in the initial radially symmetrical structure. This leads to ubiquitous activation of WNT signaling and expression of the primitive-streak marker *T/Bra* across the aggregate, resulting in a uniform induction of mesodermal differentiation. This is in marked contrast to the mouse embryo, where the WNT signaling pathway is initially activated in a highly localized manner at the primitive streak on the posterior domain, and is strictly regulated by the secretion of WNT antagonists from the anterior visceral endoderm (Arnold & Robertson 2009). The resulting signal gradient thus protects the anterior epiblast from the “posteriorizing” signals, maintaining it in an uncommitted state (Kimura et al. 2000). In fact, mutated mouse embryos with increased WNT activity upregulate the genes of the posterior mesoderm in the anterior domain (Osteil et al. 2019), which leads to a doubling of the posterior axis (Merrill et al. 2004) and to the failure of anterior brain formation (Mukhopadhyay et al. 2001; Lewis et al. 2008; Fossat et al. 2011).

Based on this understanding, we postulated that the absence of anterior neural tissue in existing gastruloids could be attributed, at least partially, to excessive WNT signaling in the early stages of culture and that, consequently, a decrease in WNT signaling levels in incipient ESC aggregates could promote the emergence of the missing anterior domains in gastruloids. In order to develop starting conditions that could trigger the formation of gastruloids independently of exogenous stimulation with the WNT agonist CHIR99021, we therefore modified the existing culture medium to obtain more physiological epiblast identity. When aggregated in high-throughput in a novel microcavity array system (Brandenberg et al. 2020), and cultured in epiblast-induction (EPI) condition in the presence of Fgf2 and activin-A, but not CHIR99021, the resulting aggregates were found to initiate *T/Bra* expression, brake symmetry and undergo axial elongation. Remarkably, in the presence of the small molecule WNT inhibitor XAV939, the EPI aggregates gave rise to gastruloids with a surprising level of antero-posterior (A-P) development, with a population of Sox1+ and Sox2+ cells in front of the extended T/Bra+ domain. Under these culture conditions, it was found that elongation and patterning efficiency was strictly dependent on initial aggregate size and WNT activity; *i.e*., smaller aggregates could not elongate and A-P patterning was abrogated without WNT inhibition. Overall, our data show the crucial role of the initial conditions, both in terms of size and cell states of the initial mESC aggregate, in promoting morphogenesis and patterning along the A-P axis. Our approach to gastruloid culture provides a simple and versatile *in vitro* tool for the study of peri-gastrulation development and especially the specification of anterior neural tissue.

## Results

### High-throughput formation of EPI aggregates in hydrogel microwells

Because the current gastruloid culture protocol is not conducive to the formation of anterior neural tissues (van den Brink et al. 2014; Turner et al. 2017; Beccari et al. 2018), likely due to premature differentiation towards mesendoderm at the expense of anterior ectoderm, here we modified the existing formulation to promote the formation of “EPI aggregates”, *i.e*. tissues that could mimic the pluripotent post-implantation epiblast that retains differentiation capacity towards both anterior ectoderm and mesendoderm lineages. We used poly(ethylene glycol) (PEG) microwells to create hundreds of aggregates of the desired size (Unzu et al. 2019; Brandenberg et al. 2020) (**Fig. 1a,b**). To avoid excessive WNT signaling, we removed CHIR99021, and we also added Activin-A (20ng/ml), Fgf2 (12ng/ml) and knockout serum replacement (1%), thus implementing a medium composition that has previously been shown to induce epiblast identity in mouse ESCs (Hayashi et al. 2011). To achieve a more homogeneous epiblast differentiation, we additionally supplemented the medium with XAV939, a small molecule inhibitor of the WNT signaling pathway (Sumi et al. 2013; Sugimoto et al. 2015).

**Figure 1:**
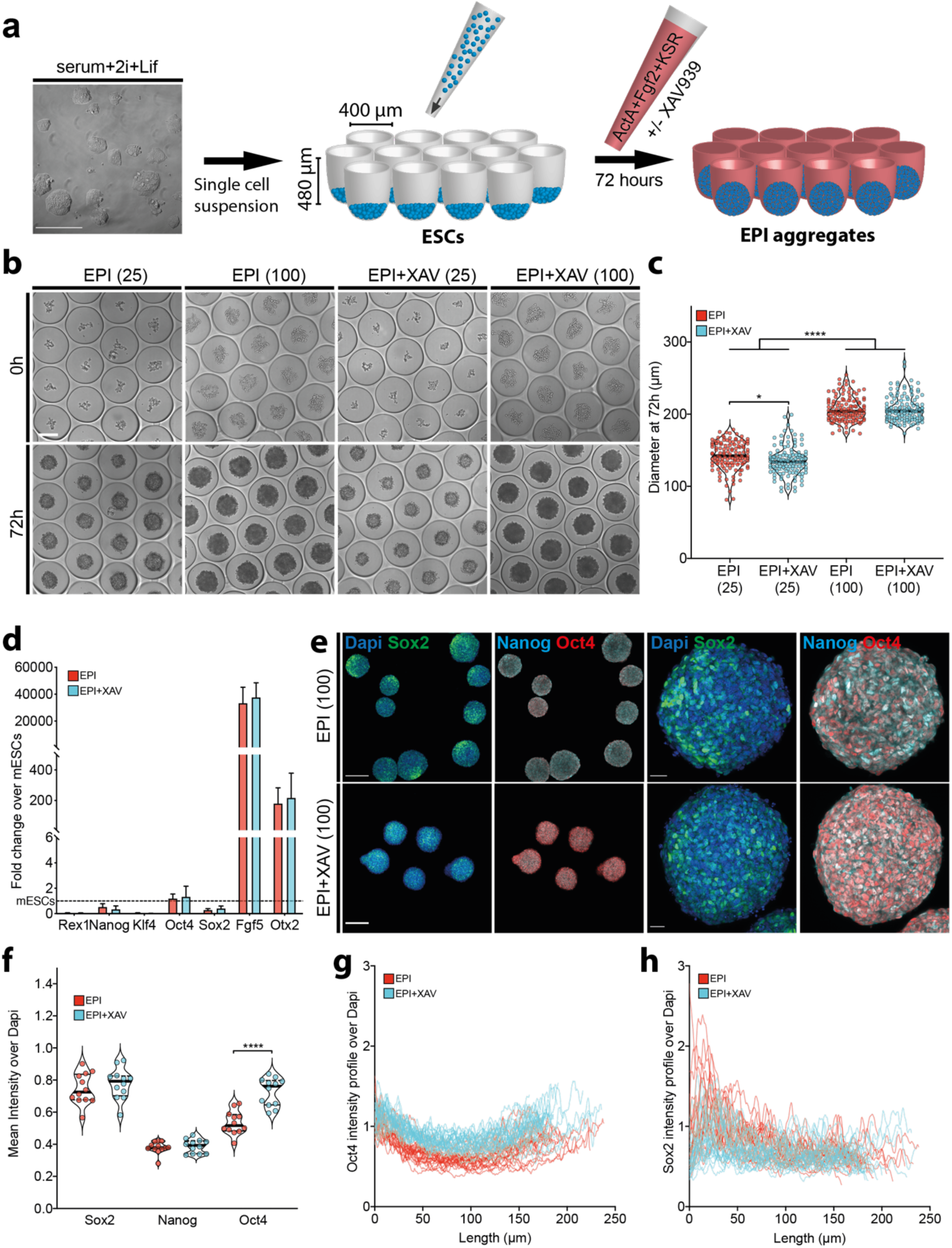
Formation of EPI and EPI+XAV aggregates on PEG microwells. **a)** Schematic showing the experimental method to generate EPI aggregates on PEG microwells from mESCs maintained in serum+2i+lif medium. **b**) Timepoint images of EPI aggregates formed from 25 mESCs/well and 100 mESCs/well at 0h (top panel) and 72h (bottom panel). **c**) Quantification of the diameter of EPI aggregates at 72h cultured in indicated conditions. Data are shown as median. **d)** qRT-PCR analysis of EPI aggregates at 72 h formed from 100 cells/well showing expression levels of pluripotency and epiblast-specific markers normalized to ESCs. Data are shown as mean with standard deviation. **e)** Confocal images showing Sox2, Nanog and Oct4 immunostainings in aggregates cultured under indicated conditions at 72h. **f)** Mean intensity value measurements of Sox2, Nanog and Oct4 normalized to Dapi intensity. Data are shown as median **g,h)** Intensity profiles of Oct4 and Sox2 normalized to Dapi in 72h aggregates cultured in indicated conditions. Lines for the intensity profile were drawn from Sox2-high pole through the midline of the aggregates. For statistical analysis, one-way ANOVA followed by Tukey multiple comparison (c,f) or two-way ANOVA followed by Sidak multiple comparison test was performed. Following P-value style was used for significance: P****<0.0001, P***<0.0002, P**<0.0021, P*<0.0332. Scale bars: 200µm.

Starting from 25 or 100 cells/well, in both EPI and EPI+XAV media, we successfully created aggregates of different sizes (**Fig. 1b**). Aggregates formed from 100 cells/well in both media had similar average diameters around 72 h (∼200µm), while aggregates formed from 25 cells/well showed a slightly reduced size when cultured in the presence of XAV939 (∼140µm vs. ∼130µm) (**Fig. 1c**).

Using ESC reporters for WNT (*TLC*:mCherry) (Ferrer-Vaquer et al. 2010; Faunes et al. 2013) and TGF-β (*AR8*:mCherry) (Serup et al. 2012), we next performed time-lapse imaging to gain insight into the dynamics of these signaling pathways during the initial culture period. As expected, WNT-positive ESCs cultured in EPI medium gradually lost reporter activity until 48h and were slightly upregulated thereafter (**Fig. S1a**). The addition of XAV939 had no effect on the initial rate of WNT downregulation, but upregulation was inhibited after 48h, resulting in significantly lower levels of WNT activity around 72h (**Fig. S1a,c**). On the contrary, TGF-β signaling was inactive in ESCs and could only be detected in aggregates after 60h. Interestingly, the addition of XAV939 led to a slightly earlier activation and higher expression of the TGF-β reporter (**Fig. S1b,c**). Of note, in both reporter systems we noticed an activation pattern running from the periphery to the center of the 400 µm microwell arrays (**Fig. S2a**). EPI aggregates formed at the periphery of the arrays up-regulated the WNT and TGF-β pathways earlier and at higher levels, suggesting a ‘border effect’ similar to the differentiation of pluripotent stem cells on circular micropatterned substrates (Warmflash et al. 2014; Etoc et al. 2016; Morgani et al. 2018). A possible explanation could be a reaction-diffusion process in which inhibitors of the WNT and TGF-β pathways were secreted from the center of the array, thereby limiting reporter activity at the periphery (Tewary et al. 2017). In support of this hypothesis, the boundary effect was eliminated when aggregates were formed in larger 800 µm wells and all aggregates uniformly activated the WNT and TGF-β reporters (**Fig. S2b**). In neither of the two tested microtiter plate arrays could we detect no *T/Bra* expression at 72h. Considering the higher number of aggregates that can be generated, we performed all the subsequent experiments on 400 µm microwell arrays.

To confirm that the aggregates have acquired epiblast identity, we next checked the expression levels of pluripotency markers by RT-PCR. Compared to ESCs, we found that the majority of pluripotency markers such as *Rex1, Klf4, Sox2 and Nanog* were downregulated in EPI aggregates, but *Oct4* expression was at similar levels. Furthermore, the expression of the epiblast-specific markers *Fgf5* and *Otx2* was highly upregulated in EPI aggregates, indicating a transition from the naïve to the primed state of pluripotency (Ghimire et al. 2018) (**Fig. 1d**). Immunostaining for SOX2, NANOG and OCT4 confirmed the expression of pluripotency factors at the protein level (**Fig. 1e**). In EPI aggregates formed in the presence of XAV939, we could demonstrate a higher and more homogeneous expression of OCT4, indicating a better maintenance of pluripotency identity with decreased WNT signaling (Kim et al. 2013) (**Fig. 1f,g**). Interestingly, we noted a somewhat polarized expression profile of SOX2 in aggregates formed in EPI condition (**Fig. 1h**). The SOX2+ pole showed an increased WNT and a decreased TGF-β activity, indicating the establishment of an anterior-posterior axis similar to that of the 48h-72h gastruloids (Turner et. al. 2017) (**Fig. S3a**). On the contrary, in the presence of XAV939, the SOX2 polarization was decreased and the EPI aggregates showed uniform activity for WNT and TGF-β reporters (**Fig. 1h, Fig. S3b**). Overall, these results suggest that ESCs could be manipulated to form epiblast-like aggregates in a scalable and reproducible manner, and that the addition of a WNT inhibitor could promote better maintenance of pluripotent epiblast identity.

### Axial morphogenesis of EPI aggregates

In conventional gastruloid culture (van den Brink et al. 2014; Turner et al. 2017) mESC aggregates assume an oval shape between 72h and 96h and continue to elongate for up to 120h. If the gastruloids are not transferred to shaking culture at this time (Beccari et al. 2018; Girgin et al. 2019), the elongation is not maintained and the gastruloids tend to acquire a rounded shape (**Fig. 2a**). To better understand the dynamics of morphogenesis over time, we performed automated image analysis by fitting a spine to the gastruloids via connecting centers of inscribed circles of different sizes (**Fig. S4a**). We calculated the elongation index by dividing the axial length by the diameter of the largest inscribed circle. This analysis showed a peak value of the elongation index at 120h, which decreased until 168h when the elongated tip collapsed (**Fig. 2b**). However, the gastruloids continued to increase in axial length up to 144h and reached over one mm, suggesting that the loss of elongated morphology was not due to growth inhibition (**Fig. S4b**).

**Figure 2:**
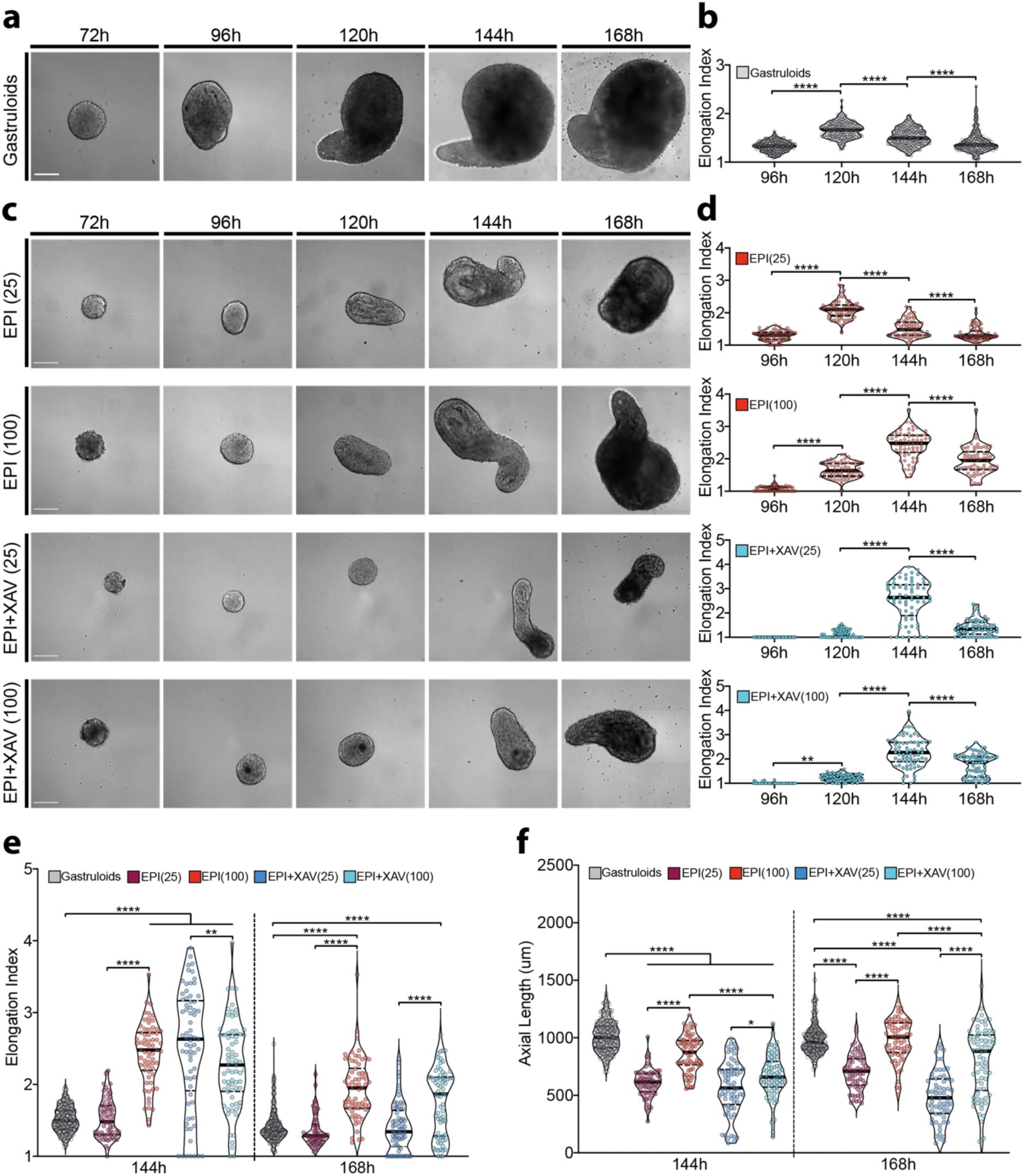
Axial morphogenesis of gastruloids, EPI and EPI+XAV aggregates. **a)** Time point images showing axial elongation dynamics of gastruloids. **b)** Quantification of the elongation index of gastruloids. Data are shown as mean with standard deviation. **c)** Time point images showing axial elongation dynamics of EPI aggregate cultured in indicated conditions. **b)** Quantification of the elongation index of EPI aggregates. Data are shown as mean with standard deviation. **e,f)** Quantification and comparison of elongation index (**e**) and axial length (**f**) of gastruloids and EPI aggregates at 144h and 168h. For statistical analysis, one-way ANOVA followed by Tukey multiple comparison test was performed. Following P-value style was used for significance: P****<0.0001, P***<0.0002, P**<0.0021, P*<0.0332. Scale bars: 200µm.

In contrast to classical gastruloids (Turner et al. 2017), when EPI aggregates were placed in low-attachment U-bottom 96-well plates after 72h, they were able to elongate autonomously and did not require external WNT stimulation (**Fig. 2c**). The elongation of EPI aggregates showed a strict dependence on the original size of the aggregates. Smaller aggregates (termed ‘EPI(25)’) assumed an ovoid shape at 96h and continued to stretch until 144h. In these structures, however, the elongation was not maintained until 168h because the aggregates gradually rounded off, similar to conventional gastruloids (**Fig. 2c,d**; **Fig. S4c**, upper row). In contrast, larger EPI aggregates (‘EPI(100)’) were more roundish at 96h and started to elongate later, but they maintained a lengthened morphology after 168h (**Fig. 2c,d**; **Fig. S4c**, second row). Surprisingly, in the presence of the WNT inhibitor XAV939 (‘EPI+XAV(25)’), smaller EPI aggregates did not extend until 120h, then started to extend massively to 144h, but could not maintain this morphology until 168h (**Fig. 2c,d**; **Fig. S4c**, third row). However, the effect of initial WNT inhibition was more subtle in larger EPI aggregates (‘EPI+XAV(100)’) as they were able to maintain an elongated morphology after 168h (**Fig. 2c,d**; **Fig. S4c**, bottom row).

The morphological analysis of the EPI aggregates grown under the different conditions confirmed that after 168h the aggregates formed from 100 cells/well were significantly more elongated than the aggregates formed from 25 cells/well (**Fig. 2e,f**). In comparison to gastruloids, larger EPI aggregates were significantly more elongated, independent of the modulation of WNT activity, but reduced in axial length (**Fig. 2e,f**). The addition of XAV939 delayed elongation and limited axial growth, but did not significantly affect the elongation index at 168h (**Fig. 2e,f**). Since focusing on elongation index alone might be misleading (**Fig. S4d**, compare EPI+XAV25 with EPI100), we selected the 100 cell/well condition, resulting in both high elongation index and axial length, for further characterization. Overall, these results show that EPI aggregates could experience axial elongation even without any exogenous stimulation. It was shown that the elongation dynamics are strictly dependent on the initial aggregate size and WNT activity.

### Anterior-Posterior patterning in EPI aggregates

Next, we used time-lapse microscopy to assess the dynamics of symmetry breaking and to monitor the emergence of *T/Bra* expression in EPI aggregates (**Fig. 3a**, upper panel). Compared to gastruloids (**Fig. S5a,b**), *T/Bra* expression was initiated 24 hours delayed in EPI aggregates. *T/Bra* expression was significantly increased after 120h (**Fig. 3b**), covering almost the entire surface of the aggregates (**Fig. 3c**). At this stage, the EPI aggregates showed a pole of T/BRA+ SOX2+ cells, which probably marked neuromesodermal progenitor (NMP) cells at the posterior tail bud of mouse embryos (Henrique et al. 2015). When EPI aggregates were further cultured until 144h, the *T/Bra* domain was restricted to the tip, surrounded by SOX2/SOX1-double positive neuronal tissue similar to the posterior structure in gastruloids (**Fig. 3e**; **Fig. S5c**).

**Figure 3:**
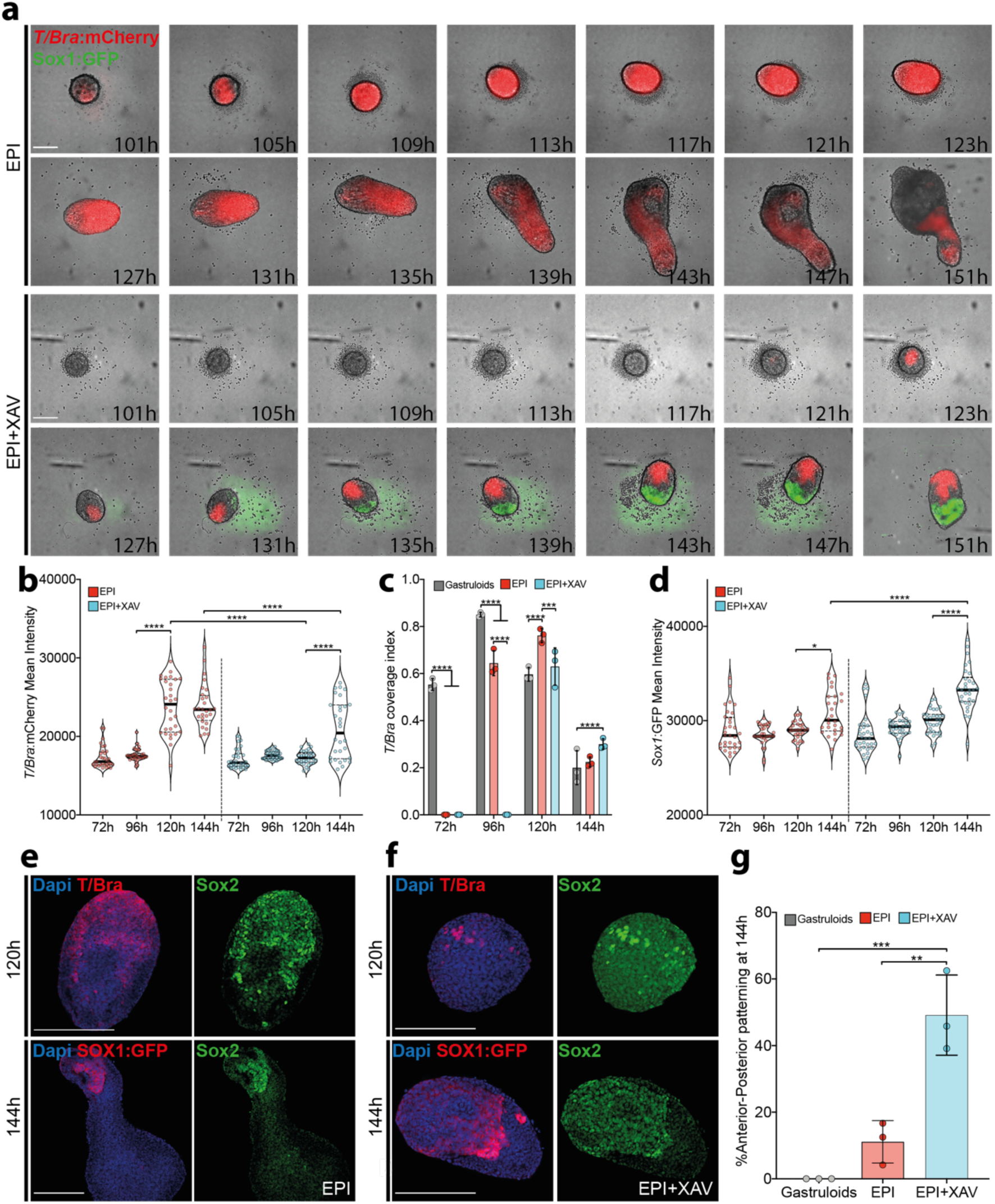
Antero-Posterior patterning in EPI and EPI+XAV aggregates at 144h. a) Time-point images showing *T/Bra* and *Sox1*expression dynamics in EPI (top panel) and EPI+XAV (bottom panel) aggregates. **b)** Quantification of mean *T/Bra* intensity in EPI and EPI+XAV aggregates until 144h. **c**) Quantification of *T/Bra* coverage index as calculated by diving the *T/Bra+* area to total aggregate area. **d**) Quantification of mean *Sox1* intensity in EPI and EPI+XAV aggregates until 144h. **e**) Representative confocal images at 120h and 144h showing posteriorly localized *T/Bra, Sox1* and *Sox2* expression in EPI aggregates. **f**) Representative confocal images at 120h and 144h showing *T/Bra* and anteriorly localized *Sox1* and *Sox2* expressions in EPI +XAV aggregates. **g**) Quantification of Anterior-Posterior patterning frequency in gastruloids, EPI and EPI+XAV aggregates described as exhibiting *T/Bra* and *Sox1* expression on the opposite sides. For statistical analysis, one-way ANOVA (**b,d,g**) or two-way ANOVA (**c**) followed by Tukey multiple comparison test was performed. Following P-value style was used for significance: P****<0.0001, P***<0.0002, P**<0.0021, P*<0.0332. Scale bars: 200µm.

EPI aggregates formed under WNT inhibition (‘EPI+XAV’) showed a delayed onset of *T/Bra* expression, starting at low levels around 120h (**Fig. 3a**, lower panel; **Fig. 3b**). In this case, it was found that the expression domain of *T/Bra* remained rather localized and did not spread across the entire aggregate (**Fig. 3c**). The *T/Bra* negative domain started to express *Sox1* (**Fig. 3a**, lower panel) and expression increased significantly up to 144h (**Fig. 3d**), indicating the formation of neural precursors on the anterior domain of the aggregates. The neural precursors could be detected already after 120h, as the EPI+XAV aggregates were uniformly SOX2-positive except for a few highly positive T/BRA/SOX2 NMP cells. At 144h, the SOX2 expression was polarized to the SOX1-positive region located in front of the elongating tip (**Fig. 3f**), an asymmetric pattern profile that we refer to as “anterior-posterior (A-P) pattern”. At 144h, the frequency of A-P patterning was highest in the presence of WNT inhibitor, with ≈ 50% of the aggregates exhibiting posterior *T/Bra* and anterior *Sox1* expression. In contrast, classical gastruloids never showed *Sox1* expression anterior to T/Bra, and aggregates formed in EPI medium (*i.e*. without XAV939) rarely exhibited the phenotype of A-P patterning (≈ 10%) (**Fig. 3g**).

To better understand the role of the key pathways involved in anterior-posterior patterning, we used the aforementioned WNT and TGF-β ESC reporter lines to track the pathway dynamics during the development of EPI aggregates. Already at 96h-101h, a low level of WNT signaling was detected, which progressively increased up to 144h and marked the extended tip (**Fig. S6a**, upper panel; **Fig. S6c**). In some cases, we could detect a smaller WNT-positive domain located anteriorly (**Fig. S6a**, white arrowhead). In contrast, TGF-β activity in EPI aggregates was gradually down-regulated and limited to the anterior domain (**Fig. S6a**, lower panel; **Fig. S6d**). Overall, the expression patterns of the WNT and TGF-β pathways in EPI aggregates were strikingly similar to those of the gastruloids (**Fig. S5d**). Aggregates formed in EPI+XAV medium showed a 12-16h delay in the activation of the WNT pathway, but continued to upregulate WNT to 144h (**Fig. S6b**, upper panel; **Fig. S6c**). Interestingly, WNT activity in these aggregates was predominantly restricted to the anterior domain (**Fig. S6b**, upper panel). Similarly, we observed a general downregulation of the TGF-β signal, but this time along the midline extending to the posterior end (**Fig. S6b**, lower panel, **Fig. S6d**).

At 168h the pattern profiles in EPI and EPI+XAV aggregates were further stabilized. Aggregates formed in the EPI medium showed extended SOX2 expression since 144h, but still marked up to the “neck” of the extended tip (**Fig. 4a**, white line). As expected, the posterior neural domain co-expressed SOX1 in the vicinity of the restricted expression of T/BRA at the tip (**Fig. 4b**). Interestingly, we were able to detect a small SOX2 expression domain at the anterior-most part in a columnar epithelium delineating the cavities (**Fig. 4a**, inlet). Immunostaining for SOX17 and OTX2 showed expression in the anterior epithelia, suggesting the formation of endodermal derivatives (Costello et al. 2015) (**Fig. 4c**). As expected, the TGF-β signaling pathway was active on the endoderm domain located anterior to the CDX2+ pole (**Fig. 4d,e**). The WNT pathway was predominantly active on the extended tail and overlapped with CDX2 expression (**Fig. 4f**). Interestingly, we could not observe the formation of large endodermal pockets in the WNT reporter cell line. In some cases, small WNT-positive domains were detected on the anterior domain, organized in epithelial rosettes that were positive for OTX2, probably with neural identity (**Fig. 4f**; white arrowheads).

**Figure 4:**
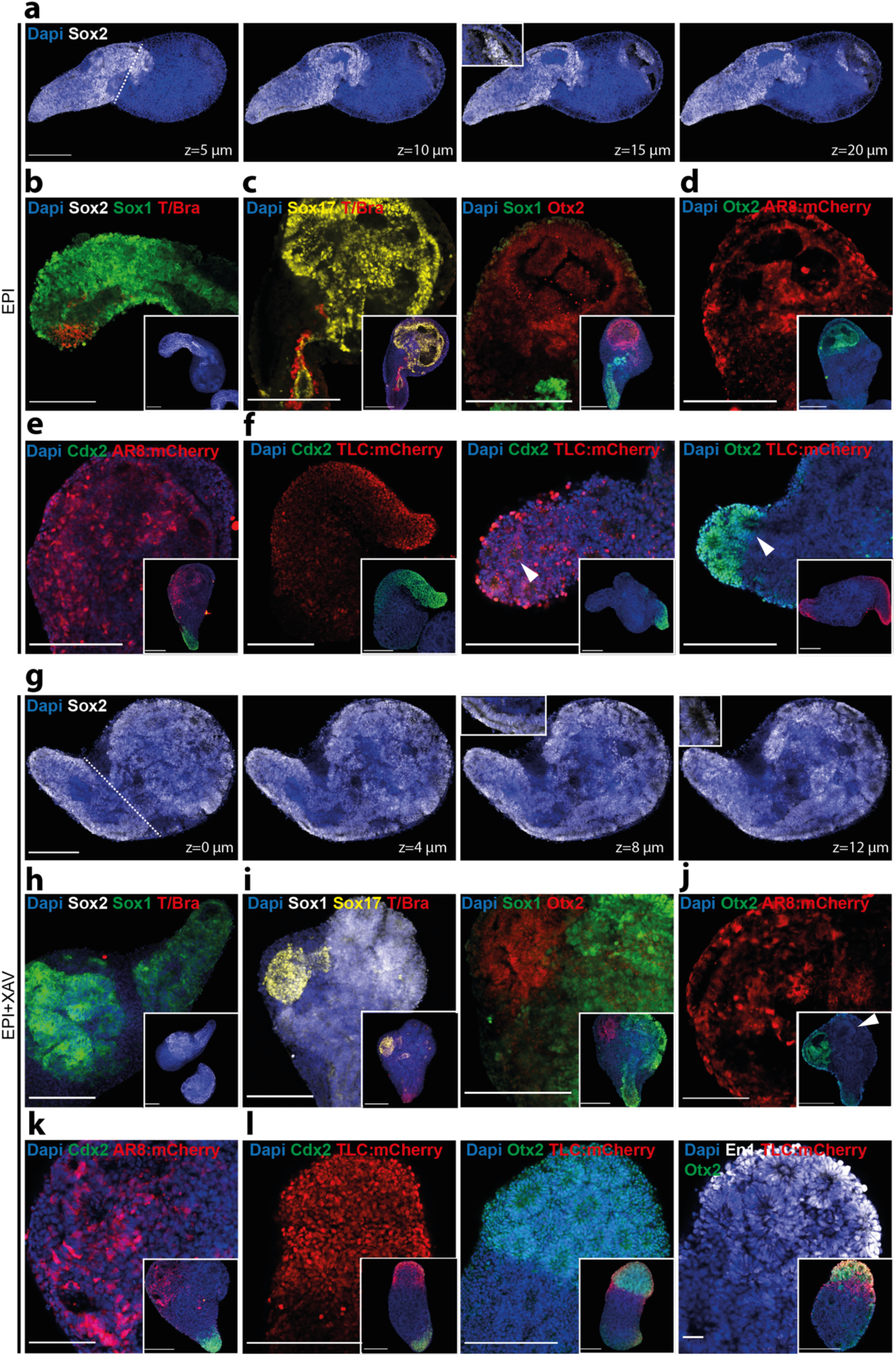
Antero-Posterior patterning in EPI and EPI+XAV aggregates at 168h. **a**) Optical sections of a representative confocal image of an EPI aggregate at 168h immunostained for *Sox2*. **b**) Confocal image showing colocalization of *Sox1* and *Sox2* expression at the posterior neural domain. **c**) Confocal images showing *Sox17* and *Otx2* expression at the anterior “endoderm” domain of EPI aggregates at 168h. **d,e**) Confocal images showing AR8:mCherry, Otx2 (**d**) and Cdx2 (**e**) expression in EPI aggregates formed from AR8:mCherry reporter line. **f**) Confocal images showing Cdx2, TLC:mCherry and Otx2 expression in EPI aggregates formed from TLC:mCherry reporter line. **g**) Optical sections of a representative confocal image of an EPI+XAV aggregate at 168h immunostained for *Sox2*. **h**) Confocal image showing colocalization of *Sox1* and *Sox2* expression at the anterior and posterior neural domain. **i**) Confocal images showing *Sox17* and *Otx2* expression at the anterior “endoderm” domain of EPI+XAV aggregates at 168h. **j,k**) Confocal images showing AR8:mCherry, Otx2 (**j**) and Cdx2 (**k**) expression in EPI+XAV aggregates formed from AR8:mCherry reporter line. **l**) Confocal images showing Cdx2, TLC:mCherry and Otx2 expression in EPI aggregates formed from TLC:mCherry reporter line. Scale bars: 200µm except (**j,k**) and (**l**, third panel) which are 100µm and 50µm, respectively.

WNT-inhibited aggregates showed anterior expression of SOX2, which marked tubular epithelia extending from the tip of the tail towards several putative neural rosettes at the anterior end (**Fig. 4g**; inlets). These rosettes were positive for SOX1 and were often located adjacent to the SOX17+ OTX2+ endodermal domain, which was much smaller than the EPI counterparts (**Fig. 4h,i**). Similarly, these small endodermal pockets were positive for TGF-β signaling, which was inactive in adjacent neuronal rosettes (**Fig. 4j**; white arrowhead). WNT signaling marked the anterior domain versus CDX2, which was organized in neural rosettes and was positive for OTX2 and EN1, indicating a possible formation of a primitive midbrain-like region in late-stage EPI+XAV aggregates (**Fig. 4k,l**).

Taken together, these results suggest that early inhibition of WNT signaling in EPI aggregates promotes the formation of anterior neural tissue. A limited posterior pattern profile in gastruloids and EPI aggregates formed in the absence of WNT inhibition suggests that anterior neural tissue precursors were probably lost in these structures and replaced by endodermal precursors instead. The addition of XAV939 could probably reverse this effect by maintaining a more homogeneous epiblast identity, as evidenced by uniform expression of pluripotency factors and delayed axis specification. In turn, these aggregates could have a much broader differentiation potential and form anterior neural and endodermal tissues composed of neural rosettes and columnar epithelia, respectively.

## Discussion

We report on a culture system for the derivation of post-implantation epiblast-like structures by aggregation and pretreatment of ESCs. We formulated a serum-free epiblast induction medium containing Activin-A (TGF-β agonist), Fgf2 (Fgf agonist) and knockout serum substitutes that promoted the acquisition of epiblast identity of ESC aggregates, followed by their spontaneous symmetry breaking and subsequent morphogenesis without any external WNT stimulation. Accordingly, mouse embryos that specifically lack Wnt3 in the posterior visceral endoderm break symmetry, initiate gastrulation and form a primitive streak, indicating an autonomous development potential of epiblast cells that is independent of an extra-embryonic WNT source (Yoon et al. 2015). It is conceivable that in our EPI aggregates a critical level of WNT signaling is reached by synergistic activities of TGF-β and Fgf pathways, as previously shown (Hayashi et al. 2011), which in turn triggers T/Bra expression (Turner et al. 2017; Berge et al. 2008). In fact, a TCF/LEF complex reporter showed endogenous WNT activity in 72h EPI aggregates, which preceded *T/Bra* expression and was further expressed until late stages. This could indicate that a WNT signal is required for the elongation of EPI aggregates, analogous to the dependence of epiblast cells on endogenous WNT signals for further development (Tortelote et al. 2013). We have shown that the inhibition of WNT signaling by XAV939 during the first 72h does not have a significant effect on elongation at later points in time, indicating a dynamic, self-organizing nature of EPI aggregates. Nevertheless, inhibition of the WNT signaling pathway at later stages or at different levels might possibly influence the dynamics of axial elongation by shifting the endogenous WNT activity threshold.

Similar to the gastruloids, EPI aggregates showed a posterior pattern with WNT+ neural tissue localized at the elongating tip (Nordström et al. 2002) and TGF-β+ endodermal tissue located at the anterior domain (Tremblay et al. 2000). This limited pattern may be due to the fact that WNT levels in early-stage gastruloids and EPI aggregates are above a threshold that would promote consistent mesendodermal conversion at the expense of anterior neural progenitors. Accordingly, Osteil and colleagues showed that embryos with increased WNT activity lose these precursors and replace them with mesoderm derivatives instead. In this work (Osteil et al. 2019), treatment with the WNT inhibitor IWP2 resulted in a more homogeneous epiblast stem cell population and increased differentiation ability compared to ectodermal derivatives. In our model, early WNT inhibition by XAV939 allowed higher OCT4 levels and a more homogeneous SOX2 expression compared to aggregates cultured only in EPI medium. This could indicate a prolonged pluripotent epiblast identity with an increased potential for differentiation into both anterior neural and posterior mesendodermal fates. As a result, we observed the occurrence of SOX1+ SOX2+ neural precursors in WNT-inhibited EPI aggregates that were adjacent to SOX17+ endodermal tissue located anterior to the *T/Bra* expressing tip. Such a pattern profile had not previously been observed in gastruloids.

Overall, our work offers a new methodology for studying early embryonic development of the mouse *in vitro* with an extended potential compared to conventional gastruloids. We believe that these second generation gastruloids could be useful to answer questions focusing on epiblast development and the formation of ectoderm and mesendoderm origin tissues. Future work is needed for a more detailed analysis of the cell types that are produced in EPI aggregates. In addition, the addition of extraembryonic cells could be a useful extension of the model, allowing to study the extent of the self-assembly properties of embryonic stem cells of the mouse.

## Materials and Methods

### Cell culture

Mouse embryonic stem cells (SBr line (*Sox1:*GFP;*T/Bra:*mCherry) (Deluz et. al. 2016), WNT line (*TLC*:mCherry) (Ferrer-Vaquer et al. 2010; Faunes et al. 2013) and TGF-β line (*AR8*:mCherry) (Serup et al. 2012)) were cultured at 37°C in 5% CO2 in medium composed of DMEM+Glutamax (#61965-026), 10% ES cell-qualified FBS (#16141-079), 1mM sodium pyruvate (#11360-070), 1x MEM non-essential aminoacids (#11140-035), 0.1mM 2-mercaptoethanol (#31350-010) and 1000u/ml Pen/Strep (#15140-122) supplemented with 3µm GSK3i (#361559), 2µm MEKi (#S1036) and 0.1µg/ml LIF (in house preparation). Cells were routinely passaged every 2-3 days by seeding 8000-9000 cells/cm2 and every 20 passages a fresh vial was thawed. Cells were tested and confirmed free of mycoplasma.

### Preparing EPI differentiation medium

N2B27 medium was prepared by 1:1 mixing of DMEM/F12+Glutamax (#31331-028) and Neurobasal (#21103-049) with the addition of 0.5x N2 supplement (#17502001), 0.5x B27 supplement (#17504001), 0.5x Glutamax (#35050-038), 1mM sodium pyruvate (#11360-070), 1x MEM non-essential aminocacids (#11140-035), 0.1mM 2-mercaptoethanol (#31350-010) and 1000u/ml Pen/Strep (#15140-122). 12ng/ml Fgf2 (#PMG0035), 20ng/ml Activin-A (#338-AC) and 1% KSR (#10828-010) were added to make final EPI differentiation medium.

### Preparing EPI aggregates on PEG microwells

Poly(ethylene glycol) (PEG) microwells with 400µm well diameter (121 wells per array) were prepared on 24-well plates as previously described (Brandenberg et. al. 2020). Microwells were equilibrated with 50µl of either EPI differentiation medium 30 minutes at 37°C. Mouse ESCs were dissociated to single cells with Accutase (#A11105-01). Cells were then centrifuged at 1000 rpm for 5 minutes and washed twice with 10 ml PBS. Cells were resuspended in EPI differentiation medium and desired cell number per well was further adjusted from the cell suspension. For example, 35µl of the 484.000 cells/ml suspension was added dropwise on microwell arrays to have 100-150 cells/well. Seeding was done at 37°C for 20 minutes. 1 ml of EPI differentiation medium with or without 10µm XAV939 (#3748) was slowly added from the side of the well. Plates were kept at 37°C in 5% CO2 for at least 72 hours before further processing.

### Transferring EPI aggregates to 96-well plates

At 72-80 hours of culture, aggregates on microwell arrays were flushed out and transferred to non-tissue culture treated 10cm plates in 10ml warm N2B27 medium with no additional factors. Single EPI aggregates were picked in 10µl and transferred to low adherent U-bottom 96 well plates (#COR-7007). 180µl of N2B27 medium was added on top. At 96, 120 and 144h hours, 150µl of medium was replaced with fresh N2B27 and EPI aggregates were kept until 168h.

### Immunostaining and confocal microscopy

EPI aggregates at different stages were washed with PBS and fixed with 4% PFA for 2 hours at 4°C. PFA was removed by three serial washes of 20 minutes at room temperature. Blocking was performed in blocking solution (PBS+10%FBS+0.3% Triton-X) for 1 hour at room temperature. Primary antibodies were incubated for at least 24 hours at 4°C in blocking solution. Next day, primary antibodies were removed by three serial washes of 20 minutes at room temperature. Secondary antibodies were incubated for 24 hours and next day EPI aggregates were washed and mounted on glass slides in mounting medium. Confocal images were taken using an LSM700 inverted (Zeiss) with EC Plan-Neofluar 10x/0.30 or Plan-Apochromat 20x/0.80 air objectives.

### Image analysis

All images were processed using algorithms developed in Image J (version 2.0.0-rc-69/1.52n). Brightfield, GFP (for *Sox1*), and mCherry (for *T/Bra, TLC*:mCherry, *AR8*:mCherry) channels were used as input. Thresholding and segmentation were performed sequentially for each channel. The coverage index was calculated by dividing area of the object identified in mCherry channel to the brightfield area. For morphology measurements, bright-field images were thresholded and segmented. Maximum inscribed circle function was used to fit circles in the identified object. Axial length was determined by connecting centers of the fit circles. Elongation index was calculated by dividing axial length to the diameter of the maximum inscribed circle.

## Acknowledgements

We thank Giuliana Rossi and Alfonso Martinez-Arias for useful feedback on the manuscript. We thank members of the Lutolf laboratory for discussions and sharing materials. We thank Romain Guiet and Olivier Burri for providing the image analysis codes, Arne Seitz and other members of Bioimaging and Optics Facility (EPFL) for microscopy support. We thank all personnel of Histology Core Facility for their technical support. This work was funded by EPFL.

## Author contribution

M.U.G. and M.P.L. conceived the study, designed experiments, analyzed data and wrote the manuscript. M.U.G. performed the experiments.

## Supplementary Figures

**Figure S1:**
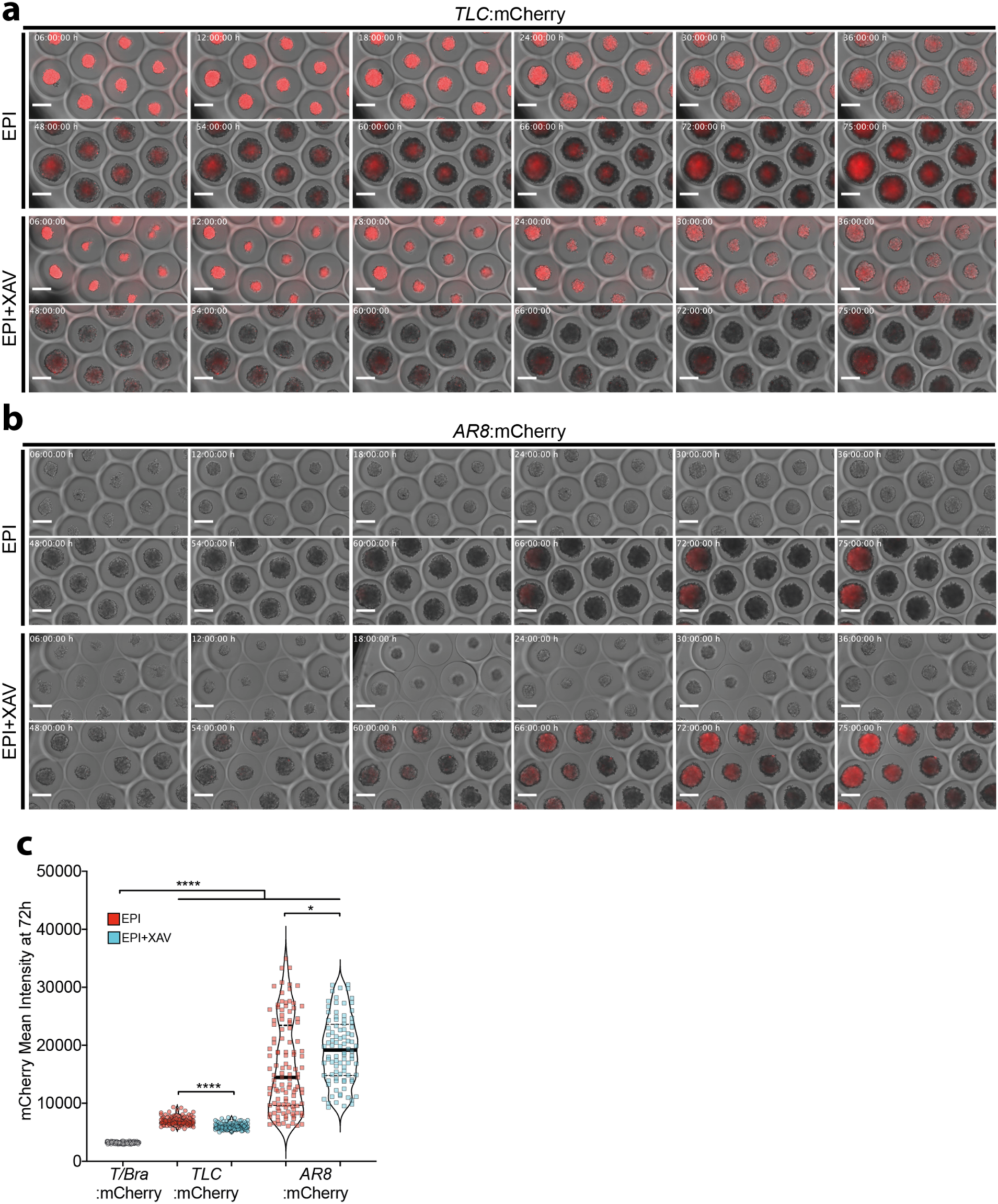
Activation of WNT and TGF-β pathways in EPI and EPI+XAV aggregates on PEG microwells. **a-b**) Timepoint images of EPI and EPI+XAV aggregates formed from 100 mESCs/well with TLC:mCherry (**a**) or AR8:mCherry (**b**) lines. **c**) Quantification of the mean mCherry intensity of aggregates at 72h formed from indicated cells lines and conditions. Data are shown as median. For statistical analysis, one-way ANOVA mixed-effects analyisis followed by Tukey multiple comparison test was performed. Following P-value style was used for significance: P****<0.0001, P***<0.0002, P**<0.0021, P*<0.0332. Scale bars: 200µm.

**Figure S2:**
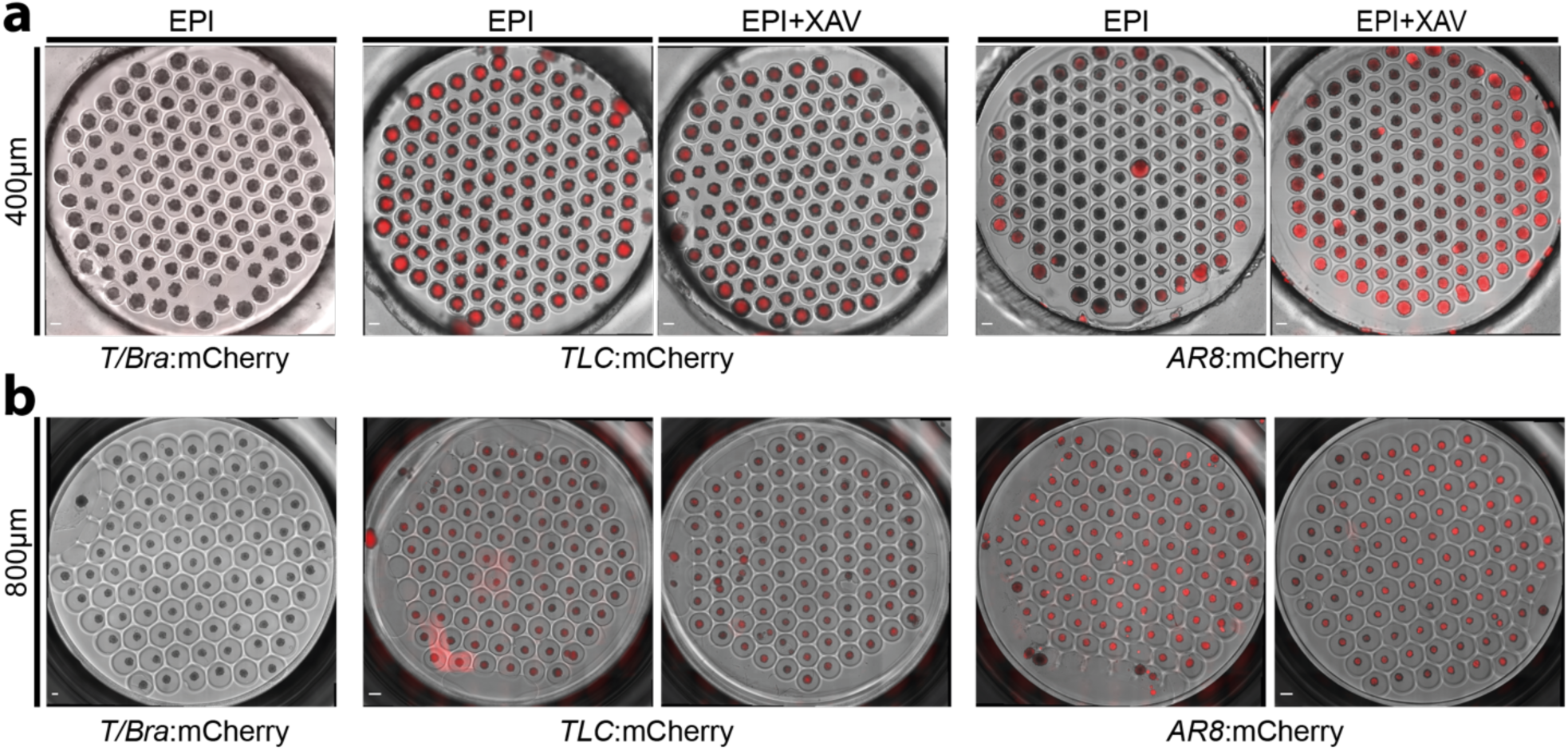
Formation of EPI aggregates on PEG microwells of different well dimeters. **a,b**) Whole array images of EPI and EPI+XAV aggregates formed from 100 mESCs/well with TLC:mCherry or AR8:mCherry lines in either 400µm (**a**) or 800 µm (**b**) microwells. Scale bars: 200µm (**a**), 400µm (**b**).

**Figure S3:**
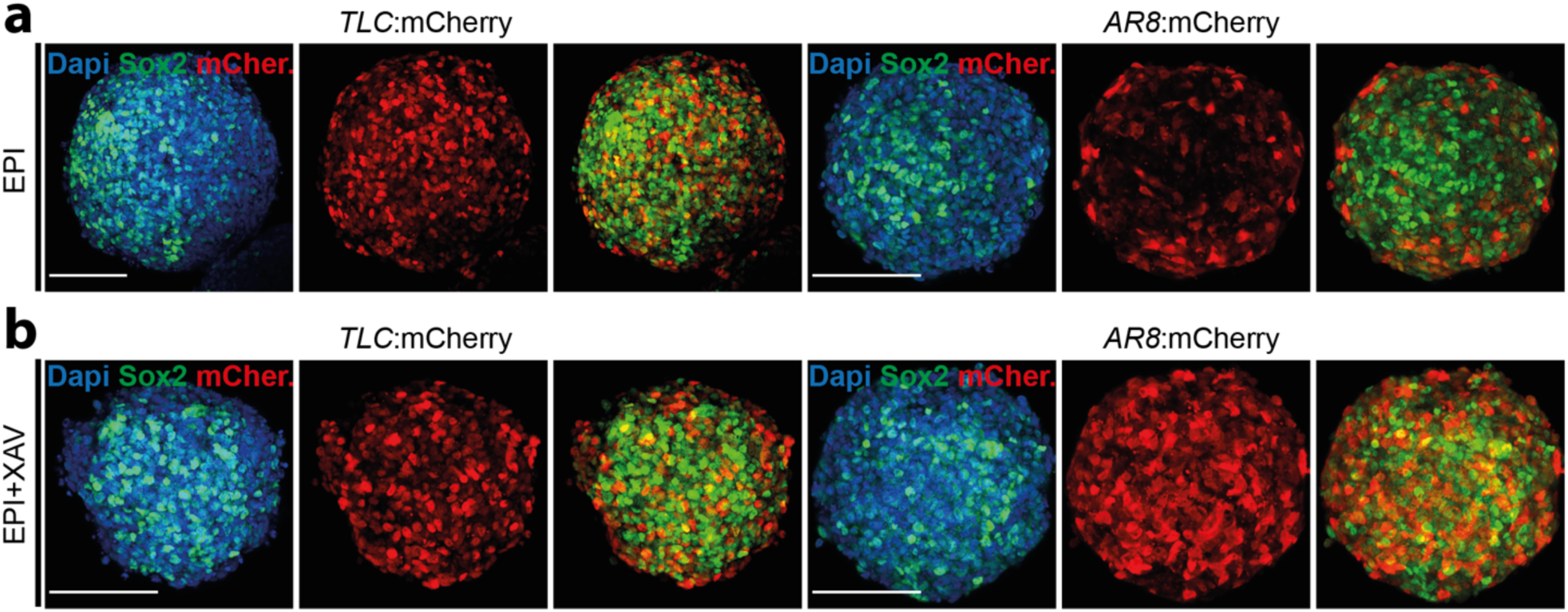
Activation of WNT and TGF-β pathways in EPI and EPI+XAV aggregates. **a,b**) Confocal images showing Sox2, TLC:mCherry and Sox2, AR8:mCherry immunostainings in aggregates cultured in EPI (**a)** or EPI+XAV (**b**) conditions at 72h. Scale bars: 100µm.

**Figure S4:**
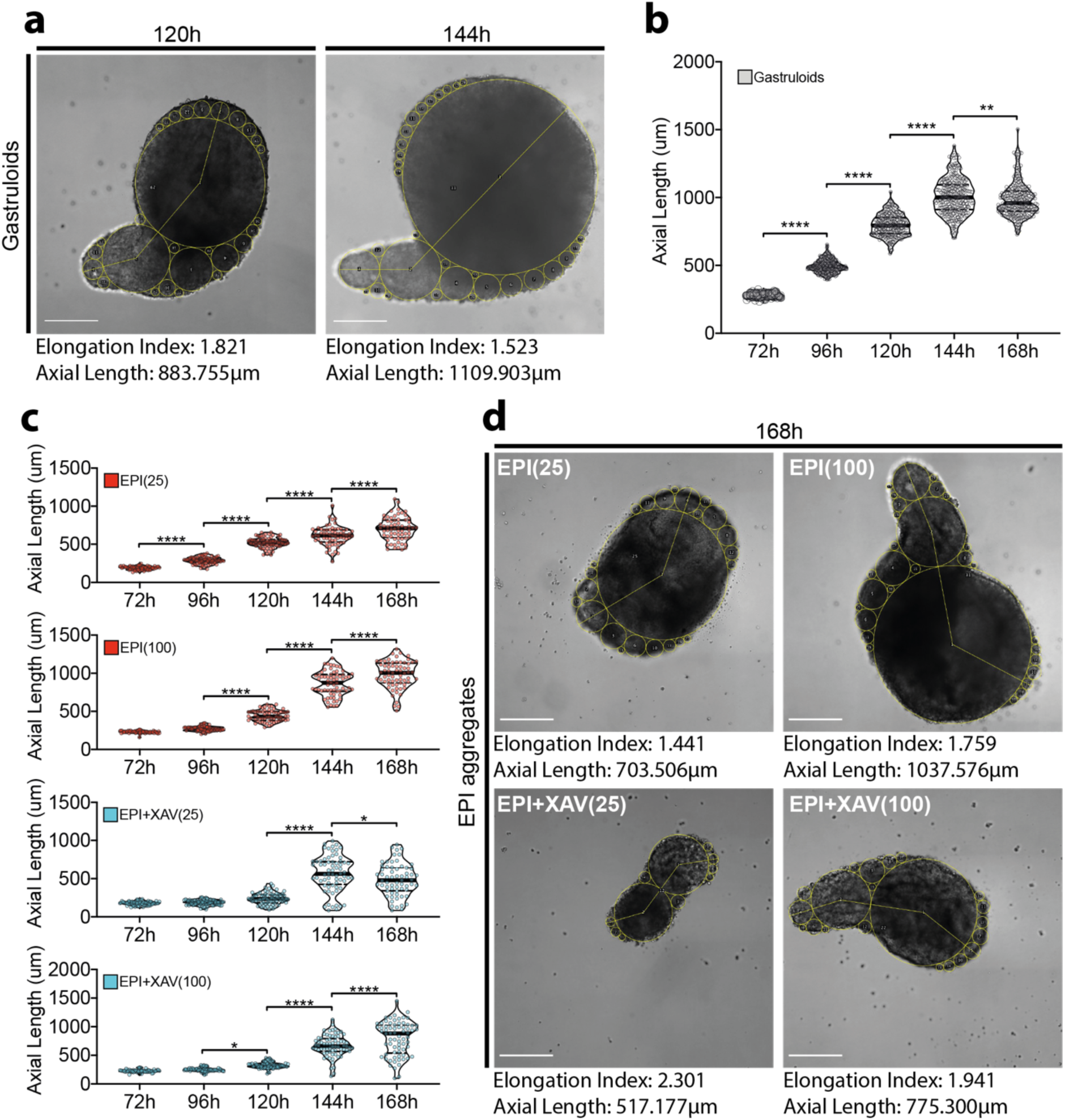
Axial morphogenesis of gastruloids, EPI and EPI+XAV aggregates. **a**) Representative images of *gastruloids* at 120h and 144h showing elongation index and axial length measurements. **b**) Quantification of axial length of *gastruloids* until 168h. **c**) Quantification of axial length of EPI aggregates until 168h at indicated conditions. **d**) Representative images of EPI aggregates at indicated conditions and their respective elongation index and axial length measurements. For statistical analysis, one-way ANOVA followed by Tukey multiple comparison test was performed. Following P-value style was used for significance: P****<0.0001, P***<0.0002, P**<0.0021, P*<0.0332. Scale bars: 200µm.

**Figure S5:**
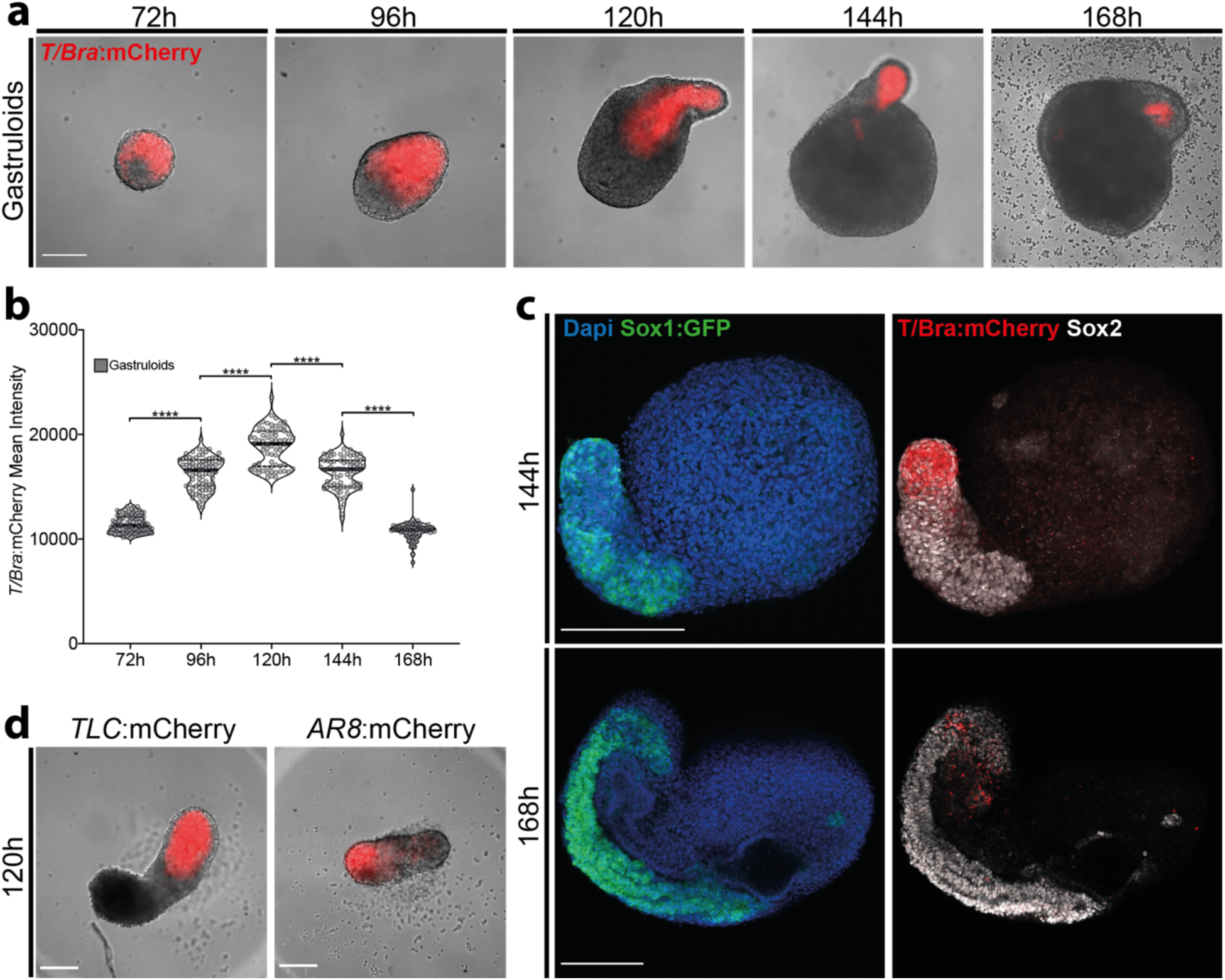
Posterior patterning in gastruloids. **a)** Time-point images showing *T/Bra* expression dynamics in *gastruloids*. **b)** Quantification of mean *T/Bra* intensity in *gastruloids* until 168h. **c**) Representative confocal images at 144h and 168h showing posteriorly localized *Sox1* and *Sox2* expression in *gastruloids*. **d)** Representative image of *gastruloids* formed from TLC:mCherry and AR8:mCherry reporter lines at 120h. For statistical analysis, one-way ANOVA followed by Tukey multiple comparison test was performed. Following P-value style was used for significance: P****<0.0001, P***<0.0002, P**<0.0021, P*<0.0332. Scale bars: 200µm.

**Figure S6:**
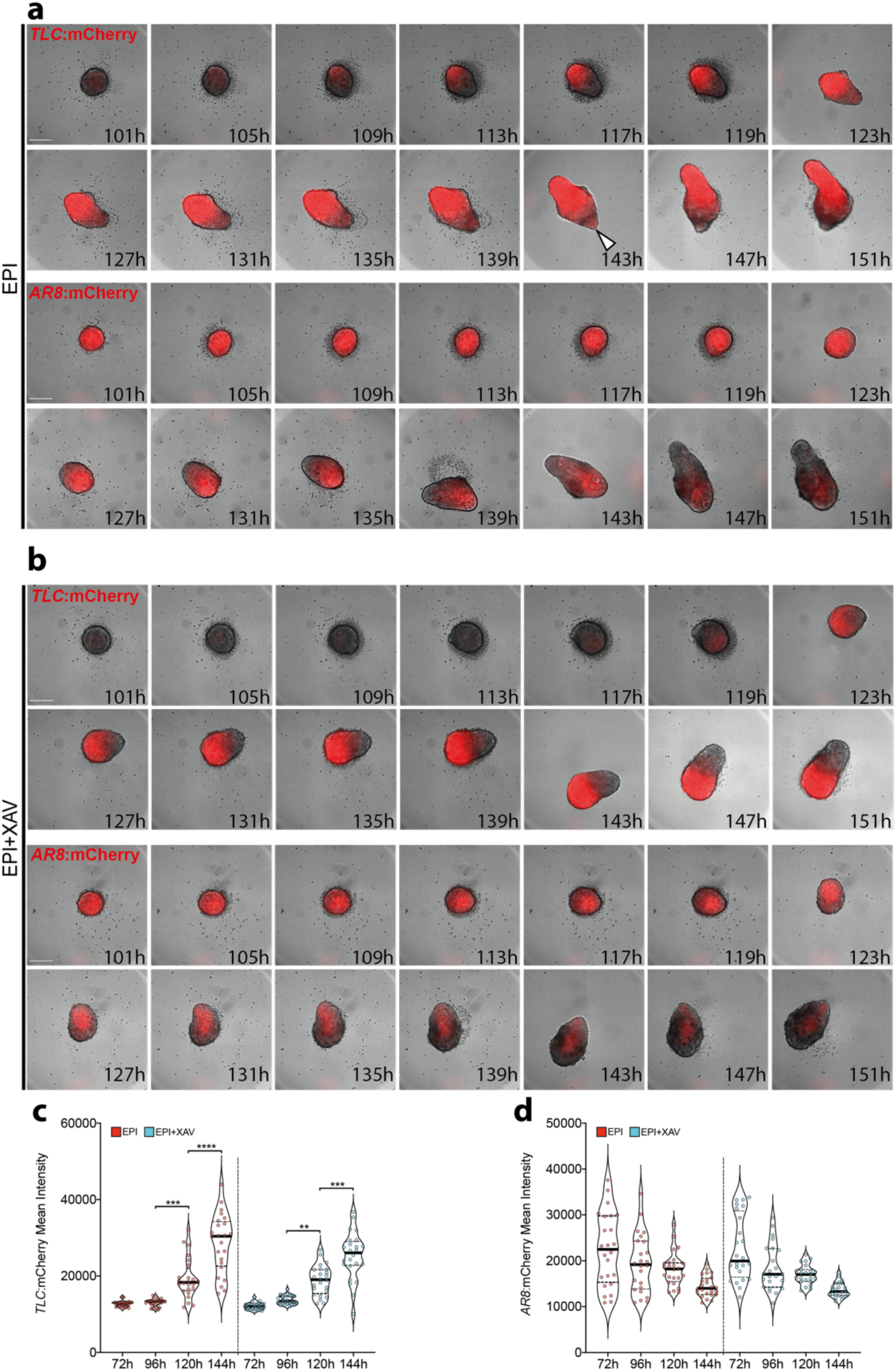
Antero-Posterior patterning in EPI and EPI+XAV aggregates formed from reporter lines. **a-b)** Time-point images showing EPI (**a**) and EPI+XAV (**b**) aggregates formed from TLC:mCherry and AR8:mCherry reporter lines. **c-d)** Quantification of mean mCherry intensity in EPI and EPI+XAV aggregates formed from TLC:mCherry (**c**) and AR8:mCherry (**d**) reporter lines. For statistical analysis, one-way ANOVA followed by Tukey multiple comparison test was performed. Following P-value style was used for significance: P****<0.0001, P***<0.0002, P**<0.0021, P*<0.0332. Scale bars: 200µm.

